# Explicit Modelling of Antibody Levels for Infectious Disease Simulations in the Context of SARS-CoV-2

**DOI:** 10.1101/2023.03.31.535072

**Authors:** Sebastian A. Müller, Sydney Paltra, Jakob Rehmann, Kai Nagel, Tim O.F. Conrad

## Abstract

Measurable levels of immunoglobulin G antibodies develop after infections with and vaccinations against SARS-CoV-2. These antibodies are temporarily dynamic; due to waning, antibody levels will drop below detection thresholds over time. As a result, epidemiological studies could underestimate population protection, given that antibodies are a marker for protective immunity.

During the COVID-19 pandemic, multiple models predicting infection dynamics were used by policymakers to plan public health policies. Explicitly integrating antibody and waning effects into the models is crucial for reliable calculations of individual infection risk. However, only few approaches have been suggested that explicitly treat these effects.

This paper presents a methodology that explicitly models antibody levels and the resulting protection against infection for individuals within an agent-based model. This approach can be integrated in general frameworks, allowing complex population studies with explicit antibody and waning effects. We demonstrate the usefulness of our model in two use cases.

## 1 Introduction

Measurable immunoglobulin G (IgG) antibodies to *Severe Acute Respiratory Syndrome Coronavirus 2* (SARS-CoV-2) antigens develop after most infections with and vaccinations against SARS-CoV-2 ([Wel+20]). Although the extent of immunity associated with different antibody titers and other immune responses is not yet fully understood, it is highly likely that an individual’s antibody level provides some information about their specific risk and severity of a future infection ([Suh+20; Kra21]). However, SARS-CoV-2 IgG antibody levels are temporally dynamic, and decrease over time if no further immunization event occurs ([Ada+20]). This waning process has been confirmed in multiple studies, showing similar effects regardless whether the immunization happened through vaccination or infection ([Che+22a; Add+22]). It has been consistently shown that the total antibody level starts declining about six weeks after the immunization event and potentially reduces by more than 50% over 10 weeks ([Shr+21; Lon+20; Seo+20]). Hence, waning is important and should be considered explicitly when modelling the antibody level.

During the COVID-19 pandemic, multiple models for projecting and predicting the spread of infections have been developed. In many countries, researchers and policy makers have been using these models to simulate and implement public health policies. From a modelling perspective, explicitly integrating antibody and waning effects into the simulation framework is crucial to allow reliable calculations of the individual risk of infection and severeness estimation. So far, only very few approaches have been suggested that explicitly treat these effects (see Sec. 4.3).

In this paper, we describe how to model antibody levels explicitly on an individual level, such that the population-wide statistics are as close to reality as possible. This approach can be integrated into general frameworks, allowing complex population studies with explicit antibody and waning effects. We demonstrate the usefulness of our model in two use cases: First, we show how to model a population, based on available data, which allows the derivation of time-dependent immunization statistics of the individuals. Second, we describe how the antibody model can be used to calculate protection levels (against infection) from virus variants for the entire population, specific sub-groups or on the individual level.

The contributions of this paper are three-fold:

1. We briefly review the current state-of-the-art literature concerning approaches for modelling individual antibody levels in epidemiological predictive models of COVID-19 (see Sec. 4.3).
2. We present an approach that allows modelling of explicit antibody levels and effects such as waning for a population or an individual, based on available data, such as vaccination and infection statistics.
3. We show how information about the antibody level from our model can be translated into an individual’s specific *protection level*. This is done explicitly for various SARS-CoV-2 virus variants (wild-type, Delta, BA.1, BA.2, etc.).

## 2 Results

The proposed model allows the calculation of an individual’s antibody level and, based on that, their protection against COVID-19 infections. By protection, we mean the reduction of the infection probability, as compared to a person without antibodies. In the context of a pandemic, this can, for example, be used to estimate the protection statistics of a country’s population. The only data needed for our model are vaccination rates and infection statistics. Both data types can usually be acquired from surveys, other models (e.g. [Mül+21]), or public sources, such as Germany’s *Robert Koch Institut* (RKI).

In the following, we show two use cases of our model (details about methods and data can be found in Sec. 4.2). All results refer to the city of Cologne in Germany.

1. We show that the model is able to calculate the number of immunization events of all population members at a given time, even for large populations such as a big city. This is the basis for more complex use cases, such as the following one. Our model is validated by the fact that its outputs closely match the observed data.
2. We show how the model can be used to gain insight into individual population groups and how they are protected against different virus strains. The model can be used as a data basis to develop strategies, such as vaccination campaigns, and can compensate for missing data.

### 2.1 Use Case 1: Population-wide immunization statistics

The model presented in this paper can be implemented as an extension of our agent-based model (ABM) ([Mül+21]). In this way, we can calculate infection dynamics that are also (but not only) dependent on immune protection. We show in the following section how the model can be used to calculate the number of immunization events (infections and vaccinations) at a given point in time. This allows us to evaluate the model in a real-world scenario by comparing it to available data. Moreover, *number of immunization events* is a relevant parameter because the strategies of policymakers often depend on what share of the population has already acquired some kind of immunity.

In Fig. 1, we show how our model results compares to observed data. Both plots show the number of exposures (vaccinations and infections) that people in different age groups had up to and including Summer 2022. The left plot contains observational data, stemming from antibody level measurements across Germany ([Lan+22]); the right plot shows our simulation results. The age group *<*18 is not shown because we do not have the observational data to compare it to. When comparing the plots, two things become clear: (1) In our model, more individuals have 4 or more exposures than shown in the observation; this implies that our model’s assumption on number of unreported cases is somewhat too high. This deviation may be due to the fact that our model’s results refer to Cologne and the data to Germany. In general, however, the deviations are small and the model results structurally fit the collected data. (2) There seem to be two groups of relatively homogeneous profiles in the model (right plot), which are not visible in the data (left plot): 18-59 and 60+. This clustering is due to the fact that the vaccination rate data for adults is only available for these two age groups. In reality, it can be assumed that, e.g., 50-year-olds have a different vaccination rate than 18-year-olds—however, we lack the data here. Although the age groups are less homogeneous in reality, it can be seen that in both the model and the data, the older groups had more frequent contact with the virus than the younger ones.

**Figure 1.**
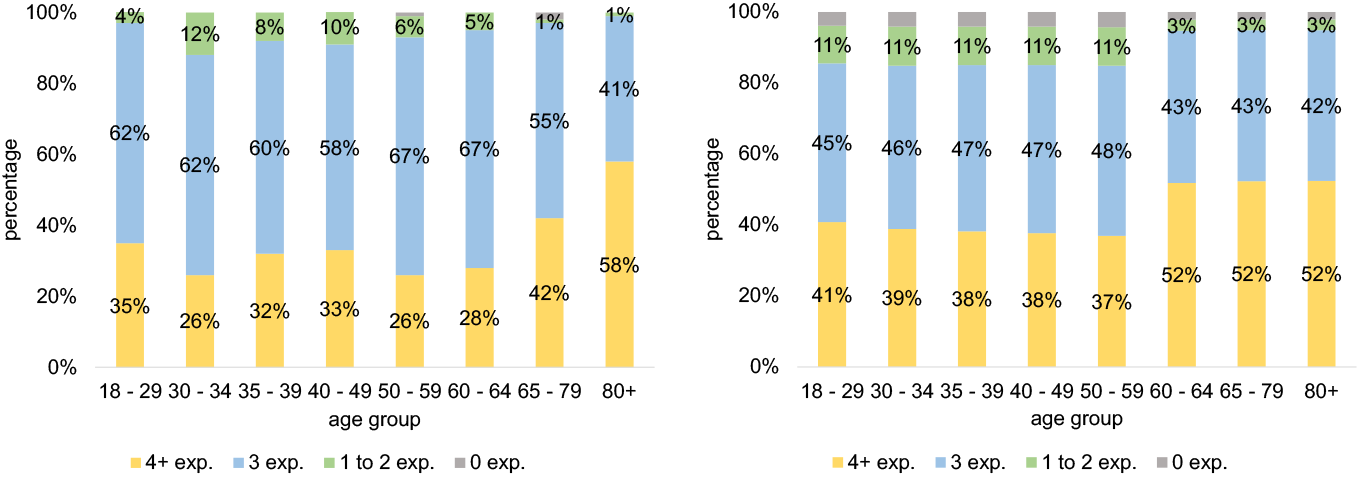
Exposures (vaccinations or infections) by end of July ’22. Left: Data from [Lan+22] for Germany. Right: Our model results for the city of Cologne.

In addition to the data shown in Figure 1, there are further surveys ([Bet+22a], [Bet+22b]) that attempted to determine what proportion of the population has had at least one infection. We only found data from summer 2022, but the available studies show results similar to our model: by then, about 35-50% of the German population had been infected with COVID-19 at least once. Since the proportion of vaccinated individuals is higher in the studies than in the general population, this value must be interpreted as a lower limit and can only be used as a rough guide. Our model calculates a value of about 50% for June 2022, which fits well to the aforementioned studies.

### 2.2 Use Case 2: Variant-specific protection of sub-groups

In general, we assume in our calculations that there is no immune protection at the beginning of the pandemic and that each infection or vaccination increases protection. The exact methodology is explained in Sec. 4.2. In the following, we depict how the population is protected against infection from different virus variants according to the model presented in this paper. Fig. 2 shows the population-wide protection against infection over time averaged over all age groups. The gray area shows that there is a large spread in immune responses; some agents are subsequently very well protected while others have almost no protection. This can partially be explained by the fact that some individuals are unvaccinated (blue dots), while others are vaccinated (red dots) or boostered (more than two vaccinations, green dots). The model results clearly show that vaccinated individuals are better protected than unvaccinated individuals, and missing vaccinations are not compensated for by infections. Thus, unvaccinated individuals do not achieve the same protection through multiple infections as vaccinated individuals.

**Figure 2.**
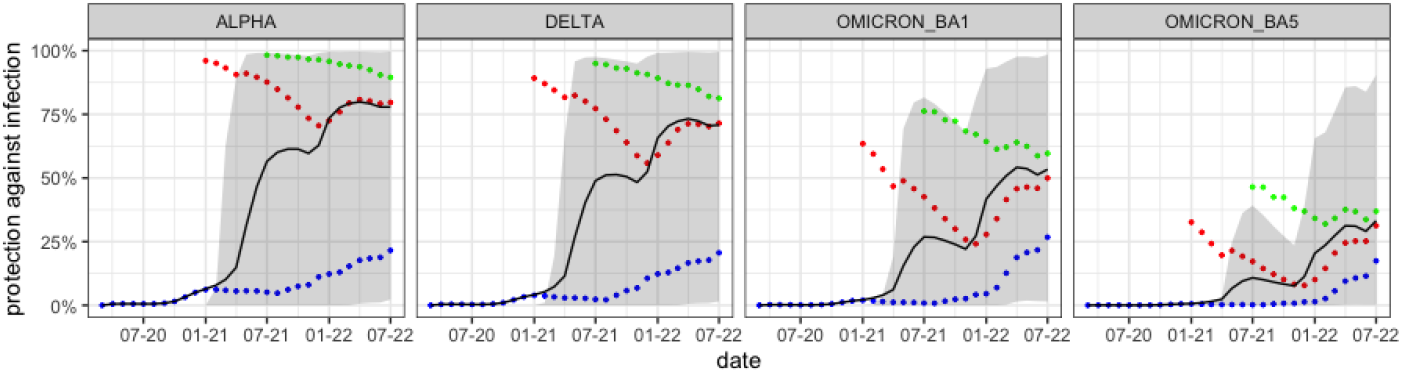
Protection against infection according to the model for different variants. The color coding is as follows: blue: unvaccinated, red: vaccinated, green: boostered, black: mean protection, gray area: 10th to 90th percentile. Reading example: for the Delta variant (2nd plot) it becomes apparent, that unvaccinated (blue) have a significantly lower protection that vaccinated (red) or boostered (green)by July 2022.

We calculate the protection for the different variants (left to right). Individuals do not have protection against any variants at the beginning of the pandemic and do not acquire significant immunity throughout 2020. This is because only a small fraction of the population was infected in 2020 and vaccinations were not yet available. Relevant immune protection is achieved by mid-2021 because vaccinations became available for the entire adult population. Beginning in July 2021, a significant decline in immune protection through waning is clearly visible. In winter 2021/2022, we see another protection increase when a third round of vaccinations (boosters) was administered to large segments of the population. In addition, the different facets of Fig. 2 show the impact of immune escape variants: in general, protection against infection with Alpha is significantly higher than against Delta, and protection against Delta is significantly higher than against either Omicron variant.

Fig. 3 shows how protection varies across age groups. It is clear that the mean protection within the age groups differs significantly. On average, children acquired less protection than adults, which can be explained by low vaccination rates in these age groups. Evidently, according to the model, the lower vaccination rates are not compensated for by infections. According to the officially reported numbers, the group of children under 5 years of age is not vaccinated at all, which results in a low level of immune protection. It also becomes apparent that the different age groups were vaccinated at different times during the vaccination campaign. The elderly over 60 were vaccinated very early, so that immune protection was also built up early. However, due to the early vaccination, there is already a significant decline in vaccination protection in the summer of 2021. In the younger adults, a similar but less pronounced effect is seen; the effect is barely visible for agents under 18. Because vaccinations were no longer administered strictly by age during the winter 2021/2022 booster campaign, it can be speculated that younger individuals had a shorter interval between 2nd and 3rd vaccination than older individuals.

**Figure 3.**
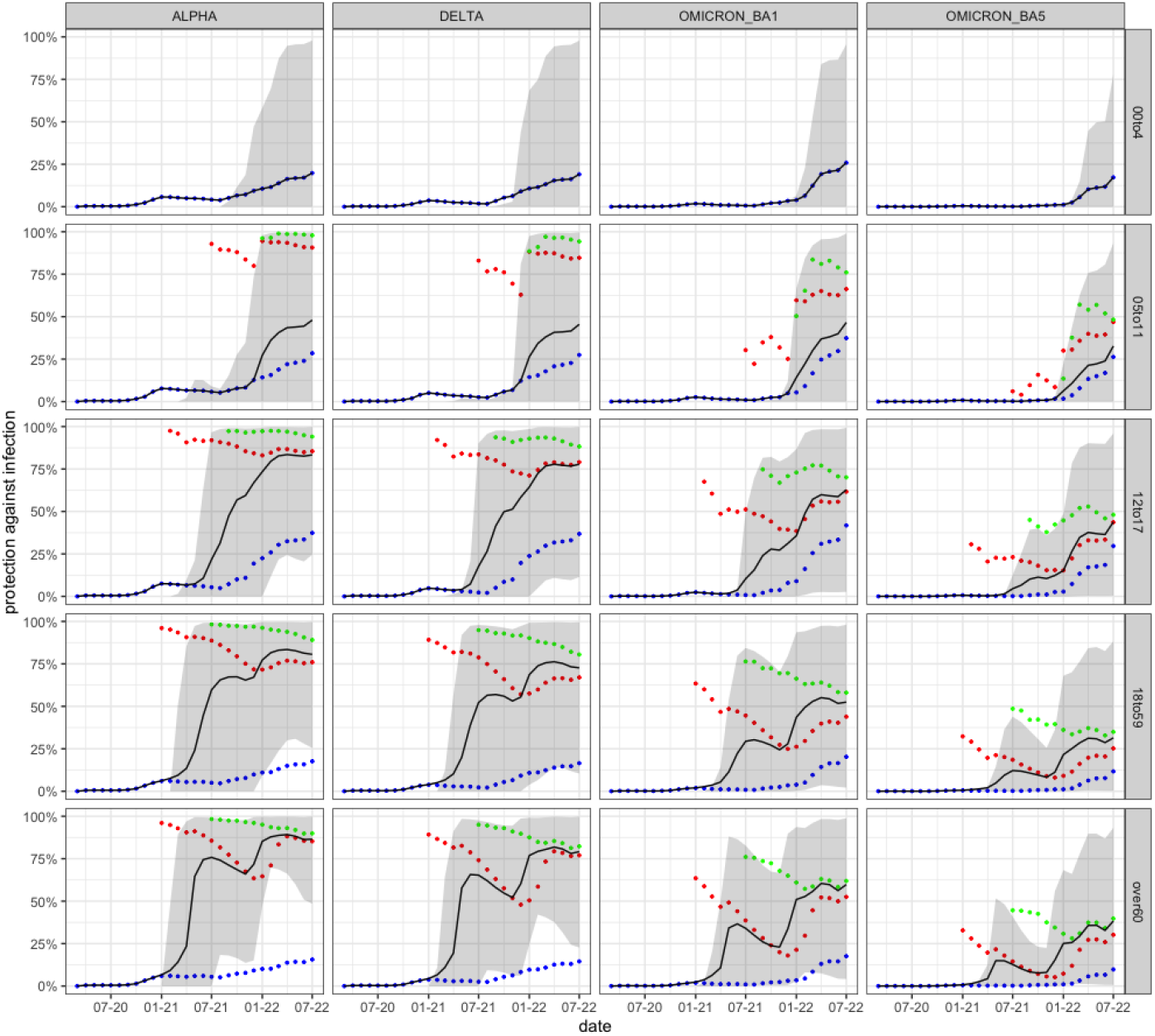
Protection against infection according to model for different variants and age groups. The color coding is as follows: blue: unvaccinated, red: vaccinated, green: boostered.

The simulation results show that protection varies significantly depending on the age group, the variant, the time point and the number of administered vaccinations. In particular, there is little protection in young children; hence, according to our model, potential vaccines for this age group could have a significant effect. In addition, it is clear that a vaccine adapted to the new variants would be helpful for all ages, since the mean protection in July of 2022 in all age groups is only about 50% or less.

### 2.3 Discussion

In the previous sections, we have shown two exemplary use cases that demonstrate the capabilities of our approach to model an individual’s antibody level and, based on that, an individual’s level of protection against infection. The data needed for our model is usually available from public sources. The presented findings for the study-area of Cologne demonstrate that the model can produce realistic results. In particular, it is possible to derive immune protection of individuals—or the entire population—as a function of time, age, vaccination status and, specifically, virus variant. It should be noted that the model is currently parameterized only for SARS-CoV-2 and its variants. It is possible, however, to adjust the model to be applicable to other infectious diseases, such as influenza.

The presented model is based on certain assumptions and simplifications. We assume, for instance, that there is a direct correlation between antibody levels and protection against infection. Moreover, we currently do not distinguish between the effects of vaccinations and infections, except for the individual’s *first* immunization event. In addition, we assume a constant antibody level half-life, regardless of whether the antibodies come from infections or vaccinations (see Sec. 4.2 for details). This can be improved in the future when more bio-medical research on SARS-CoV-2 becomes available.

## 3 Summary & Conclusion

We have presented an approach on how to model the variant-specific neutralizing effect of antibodies and how to convert it into a protection against infection. The presented use cases demonstrate that the model produces valid results that match the observed historical data in Germany very well. Further, we have shown how this approach can be used within an agent-based modelling framework to allow computation of infection dynamics. In the (current) situation of high population immunity, considering immune protection is essential for achieving realistic simulation results.

Our simulation results show that in summer 2022 there was still a significant difference in immune protection between unvaccinated and vaccinated individuals. According to the model, the lack of vaccination is not fully compensated by infections. This effect also becomes clear when looking at the age groups: according to the model, children have a significantly lower protection against infections than adults. In addition, the model allows quantification of the protection against the immune escape variants. These results suggest that the protection against the Omicron variants is significantly lower than against the original (wild-type) variant. This matches the available data. The necessary model parameters have either been taken from the available literature or are based on calibration to available data. This process necessarily includes modelling choices. Given the solid agreement between our model results and the available data, we are confident that sensible parameters and fitting parameter values have been identified. This is also confirmed by simulation results that have been achieved by using the presented antibody model in conjunction with our own agent-based model (see, for example, [Mül+22a] and [Mül+22b]). These results demonstrate once more that the agent-based model with the presented antibody-model extension is able to soundly replicate many important parameters, such as case numbers, R-values, and hospitalizations. This is a significant improvement based on the explicit antibody model for each agent.

To the best of our knowledge—and based on the our literature review—no other currently available model allows both (1) the integration of antibody levels as a proxy for protection against infection and (2) the modelling of individual immunization histories. While a small number of models implemented one of these, we haven’t encountered any that implement both. In consequence, our approach could help others to integrate any permutation of immunization events, as well as waning, into their COVID-19 models.

### 3.1 Limitations of the study

The model, as it is, works well for the presented use cases. However, limitations and possible improvements exist, and will be briefly described in the following.

On the technical side, the model is currently designed to be compatible with ABMs, such as ours [Mül+21]. However, adaption to other models types would be straightforward, since the entire source code [Rak+22] and data [Bal+21] is publicly available. From a data perspective, the presented approach is based on the immunization histories of individuals, which we generated based on real data in the presented use cases. This works well for larger populations, as demonstrated, but other approaches might be chosen for different contexts. Further, additional demographic data such as home location, income, or number of household members could be integrated into the model as well. We did not evaluate the effect of this additional information, but we intend to in a future version. The main limitation of our model from a bio-medical standpoint is that not all necessary model parameters can be derived from the literature; the reason is that the studies simply haven’t yet published. However, our approach for parameter estimation seems to work well for the presented use cases.

## 4 STAR Methods

### 4.1 Resource availability

#### Lead Contact

Further information and requests for source code and data should be directed to and will be fulfilled by the lead contact, Sebastian Müller.

#### Materials availability

This study did not generate new materials.

### 4.2 Method Details

In the following, we describe how our model computes antibody levels and how it calculates the ensuing protection against infection with SARS-CoV-2. We explain how the model parameters were chosen using both available data and our own calibration, which was necessary to fill data gaps.

The model is composed of two layers: 1) Modelling the antibody level, based on real-world measurements of antibody titers. 2) Translation of the antibody level into protection against infection, as this is the relevant parameter to calculate the infection probability.

The model is designed as an extension to our agent-based model presented in [Mül+21]. The protection is integrated as an additional parameter into the infection model, described on page 5 of that paper. The principle is simple: the higher the antibody level, the higher the protection. And the higher the protection, the lower the probability of infection, given contact with an infectious agent.

#### 4.2.1 Background

We used the models of [Coh+21] and [Cro+22], with details for the latter in [Kho+21], as starting points for the process of integrating antibodies into our agent-based model. They both postulate a logistic model of type

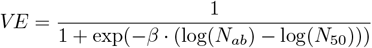

for vaccine effectiveness, where *N*_*ab*_ is the measured antibody level, *N*_50_ is the antibody level at which *VE* is 50%, and *β* determines the slope at *N*_*ab*_ = *N*_50_. Translated into relative risk, which we here call *immFac*, this can be rearranged to

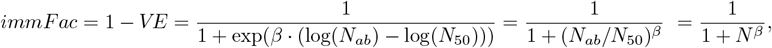

where *N* as a strain-specific relative antibody level and is defined as *N* := *N*_*ab*_*/N*_50_ (see Sec. 4.2.3 for a detailed explanation). *N* is unit-less and would need to be multiplied with *N*_50_ to be expressed in laboratory units. Note that *N* is time-dependent, as antibodies decrease over time and increase, when an infection or vaccination occurs (see Sec. 4.2.3). The value for *β* is chosen through calibration (see also Sec. 4.2.4). The equation shows that a relative antibody level of 0 leads to an immunity factor of 1, i.e. a *V E* of 0%. An antibody level of 1 leads to an immunity factor of 0.5, i.e. a *V E* of 50%. An antibody level above 1 corresponds to an immunity factor below 0.5, i.e. a *V E* higher than 50%.

#### 4.2.2. Integration with a dose-response model

Our agent-based model [Mül+21] uses the following well-established dose-response model to calculate the probability of infection ([Wat+10; SC10; Kri+21]):

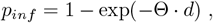

where *d* is the viral dose, and Θ is a calibration parameter, which depends on the transmissibility of the virus under consideration.

The open question was how to include *immFac* into the above dose-response infection model; since most simulations use a compartmental approach, they do not need to resolve this issue. A possible form, 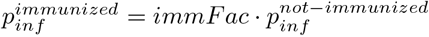, would imply full protection for people with high antibody levels, even in virus rich environments ([Che+21]). This does not seem plausible given that the virus eventually overcomes the antibodies if the ratio of virus to antibodies is large enough.

As a consequence, we put *immFac* into the exponent, as such:

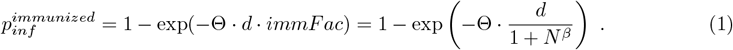

Note that this has the consequence that in a virus-limited environment, where dose *d* is small, *immFac* becomes a risk reduction for situations in virus limited environments:

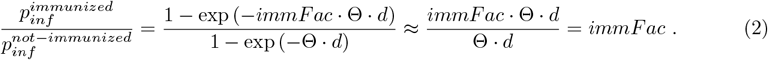

This linear approximation in a virus-limited environment follows from 1 − *exp*(− *x*) ≈ *x* when *x* ≥ 0 and sufficiently small.

That is, a model that was originally developed for a macroscopic situation is now used at a more microscopic level. The *epidemiological* risk reduction would come out as an average over many exposures with different values of *d*.

Eq. (1) shows how antibodies reduce the likelihood of becoming infected (reduced susceptibility).

However, we also included the fact that agents with antibodies have reduced probability to pass on on the virus (reduced infectivity). Thus, when an unvaccinated agent has contact with a vaccinated agent, the unvaccinated agent indirectly benefits from the vaccinated agent’s antibodies because the probability of infection is reduced. If both agents are vaccinated, the probability is further reduced. This is in accordance with findings by [Eyr+22]. In our model, the infectivity is reduced according to the same principle as explained above, but to a lesser extent. The effect of the antibodies on the infectivity is 25% of the effect they have on the susceptibility. Thus, if an agent has a 50% reduced probability of infection due to their antibodies, the probability of transmission is reduced by only 12.5%.

#### 4.2.3. Modelling the antibody level

In the next step, the relative antibody levels (*N* in Eq. (1)) are modelled. For every simulated day and agent, the model updates the agent’s relative antibody level with respect to each SARS-CoV-2 strain. A relative antibody level of 0 corresponds to no protection, while a relative antibody level of 1 corresponds to 50% protection (see Sec. 4.2.1 for details). At the beginning of the simulation, all agents are initialised with a relative antibody level of 0. Immunization events (vaccinations and infections) increase an agent’s relative antibodies. On days on which no immunization event occurs, the antibody levels follow an exponential decay curve, as shown in Eq. (3):

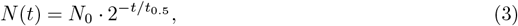

where *N* (*t*) is the antibody level on day *t* after the most recent immunization event, *N*_0_ is the antibody level immediately after the most recent immunization event and *t*_0.5_ is the half-life, which is 60 days in this case. The value of 60 days is rather on the lower end of what can be found in the literature ([Cro+22], [Yam+22], [Gil+22]). However, the value is the result of our calibration: at 60 days, our model best matches the literature in terms of the declining level of protection against infection acquired through vaccination (due to waning antibody levels). See also Sec. 4.2.4 for more details.

The general principle of the model is shown as an example in Fig. 4. The left figure shows how the antibody level of an example agent develops over time. The spikes in relative antibodies correspond, in this illustrative example case, to a SARS-CoV-2 infection, an mRNA vaccination, and an infection with the Delta variant. On days without an immunization event, the waning becomes apparent. In addition, it becomes clear that we distinguish between the different virus variants. As a result, this means that agents are less protected against the immune escape variants after vaccination. The right plot shows how we translate the antibodies into protection. See Sec. 4.2.1 and Sec. 4.2.2 for details.

**Figure 4.**
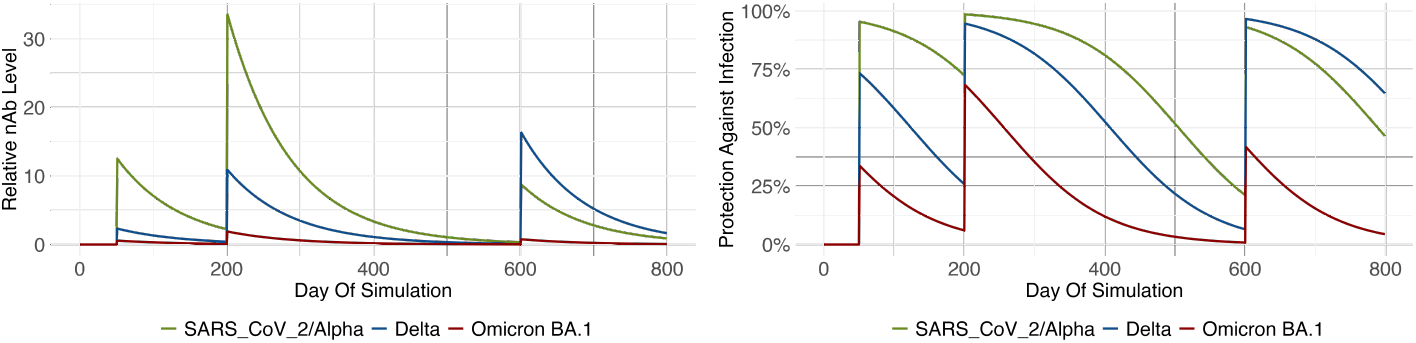
Exemplary immunization history. The agent gets infected with the wild-type on day 50, receives the mRNA vaccine on day 200 and gets infected with the Delta variant on day 600. Left: Neutralizing antibody levels, Right: Resulting protection against infection (Protection is computed as follows: 1 *− immFac* = *VE*).

##### Initial Immunization

As noted above, we assume that initially (at the beginning of the pandemic) all agents have an antibody level of 0. The first immunization event generates a strain-dependent initial antibody level, which is shown in Tab. 1. The agent’s antibodies have varying neutralizing effects against different SARS-CoV-2 strains. Thus, we model that an agent has a different relative antibody level per strain. As shown in Tab. 1, an infection with Delta provides more relative antibodies against a reinfection with Delta than against an infection with an Omicron variant. Similarly, the vaccinations were designed to protect against the wild-type and Alpha variants; thus, the vaccinations provide more relative antibodies against these strains than for later variants.

Tab. 1 is based on studies that examined protection against (symptomatic) infection and on various studies that measured antibody titers after vaccination or infection ([Rös+22b], [Rös+22a]). Here, protection obtained through vaccination with the mRNA vaccines developed by Moderna (mRNA-1273), and by BioNTech-Pfizer (BNT162b2) are summarized under ‘mRNA’, while the vector vaccines developed by AstraZeneca (ChAdOx1-S) and Johnson & Johnson (Ad26.COV2.S) are summarized under ‘vector’. In consequence, we do not distinguish between vaccine brands, but only between vaccine types.

The starting point for Tab. 1 was protection after vaccination with an mRNA vaccine against the wild-type, the Alpha, the Delta and the Omicron BA.1 variant (marked with *⋆* in Tab. 1). For these cases, studies that assess vaccine effectiveness over time are available ([NBN22], [And+22; UKH22a; UKH22b; Che+22b]). To match these studies, the corresponding initial antibody values in Tab. 1 were calibrated. In the same step, the half-life of 60 days from Eq. (3) was estimated (for the calibration process see Sec. 4.2.4 and for the conversion between vaccine effectiveness and neutralizing antibodies see Eq. (4)).

In the next step, we used measurements from [Rös+22b] and [Rös+22a] to populate the other entries. For example, the second row of Tab. 1 represents the relative antibodies versus various strains resulting from a vector vaccination. For Alpha, [Rös+22b] measure a neutralizing effect of approximately 700 after mRNA vaccination and approximately 210 after vector vaccination (we obtained these values from Figure 1 in [Rös+22b]). We used this ratio to calculate the relative antibodies against Alpha after vector vaccination: 29.2 *·* 210/700 = 8.76. The remaining entries in the table were filled following the same logic.

The measurements by [Rös+22b; Rös+22a] and others show that there is virtually no neutralizing effect if the initial immunization event is an Omicron infection, so we assume a very low value (0.01) here. We do not use 0, as it is to be expected that at least a small protection is present in the case of repeated infections.

For Omicron BA.2 and BA.5 we did not have accurate measurements at the time of the study, so we calibrated the immune escape by using our agent-based model. Here, we take the values for BA.1 from Tab. 1 and divide them by a factor. The factor was calibrated so that our model correctly replicates the infection dynamics, in particular the initial growth of BA.2 and BA.5, respectively.

**Table 1:** Initial relative antibodies per variant after certain immunization events. Based on calibration (*⋆*, see Sec. 4.2.4 for details) and [Rös+22b; Rös+22a]. ‘mRNA’ means a vaccination with an mRNA vaccine (either mRNA-1273 or BNT162b2), ‘vector’ means a vaccination with a vector vaccine (either ChAdOx1-S or Ad26.COV2.S), and ‘SARS CoV 2/Alpha/Delta/BA.1’ means an infection with the ‘SARS_CoV_2/Alpha/Delta/Omicron BA.1’ variant.

|  | initial relative antibodies against |  |  |  |
| --- | --- | --- | --- | --- |
| immunization event | SARS-CoV_2 | Alpha | Delta | BA.1 |
| mRNA | 29.20* | 29.20* | 10.90* | 1.90* |
| vector | 8.76 | 8.76 | 5.45 | 0.38 |
| SARS-CoV_2 | 12.5 | 12.5 | 2.33 | 0.57 |
| Alpha | 12.5 | 12.5 | 2.33 | 0.57 |
| Delta | 8.76 | 8.76 | 16.4 | 0.76 |
| BA.1 | 0.01 | 0.01 | 0.03 | 0.21 |

##### Agent heterogeneity

To account for the fact that immune response towards vaccinations or infections varies across the population, we assign an *immuneResponseMultiplier* to each agent. The lowest possible *immuneResponseMultiplier* is 0.1, which is an attempt to adequately depict the immunocompromised population; the maximum multiplier is 10.0. Tab. 1 presents the initial antibodies for an individual with an average response to immunization events (*immuneResponseMultiplier* = 1.0); for low and high responders, the antibodies shown in the table are multiplied by an agent’s *immuneResponseMultiplier* to calculate the antibodies gained in response to an immunization event. A log-normal distribution of *immuneResponseMultiplier* with a median of 1.0 is applied to the population.

##### Subsequent Immunizations

If the agent is subject to an additional immunization event, their antibody levels will be multiplied by a factor of 15 across the board for vaccinations and infections ([Atm+22]). The maximum antibody level that an agent can have is 150 (which corresponds to a protection of nearly 100%). If multiplication by 15 still leads to a lower protection than indicated in Tab. 1, then the value from Tab. 1 is used. This means that, at minimum, the initial antibody level from Tab. 1 is always reached.

#### 4.2.4 Calibration

As not all necessary parameters were available in the literature when we built this model, some had to be estimated. These estimations were based on studies on vaccine effectiveness and Eq. (1). The relative risk of an immunized individual vs. a non-immunized individual given dose *d* is 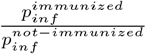. Vaccine effectiveness is defined as one minus this relative risk:

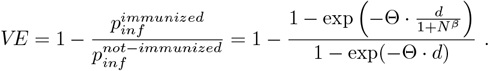

This depends on the dose *d*; for example, for *d* → 0 one obtains 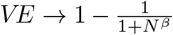, while for *d* → ∞ one obtains *VE* → 0. That is, according to the model, immunity can be overcome by a sufficiently high dose. This is similar to the distinction between virus-rich and virus-limited environments, where protection measures such as masks only make a difference in virus-limited environments [Che+21].

We performed the calibration for the exemplary value of Θ *· d* = 0.001. We also tested the calibration for values other than 0.001 and obtained very similar results, so that the value can be understood as a placeholder. In the model of [Mül+21], a value of 0.001 corresponds to contact with a contagious person for about 1000 *sec* without protection (e.g, masks) in a room of 20 *m*^2^.^1^

From the above, one obtains the following equation that can be used to convert *immFac* (and therefore the antibody level) to vaccine effectiveness against infection and vice versa:

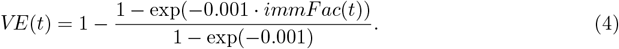

By use of the above equation we performed parameter estimations for:

- *t*_0.5_ in Eq. (3)
- Entries marked with *⋆* in Tab. 1
- *β* in Eq. (1)

To this end, for the wild-type/Alpha, the Delta, and the Omicron BA.1 variants, Eq. (4) was fitted to data points taken from studies ([NBN22; And+22; Che+22b; UKH22a; UKH22b]) by minimizing the mean squared error. For this, we used R (version 4.1.1) and the optim function from the stats package ([R C21]). Optim can be used for general purpose optimization as it is based on Nelder–Mead, quasi-Newton and conjugate-gradient algorithms. The results can be seen in Fig. 5, where the dots are values taken from the studies mentioned above, and the lines show the vaccine effectiveness *VE*(*t*) in our model when using the calibrated values. The increased vaccine protection after 210 days is related to booster vaccinations.

**Figure 5.**
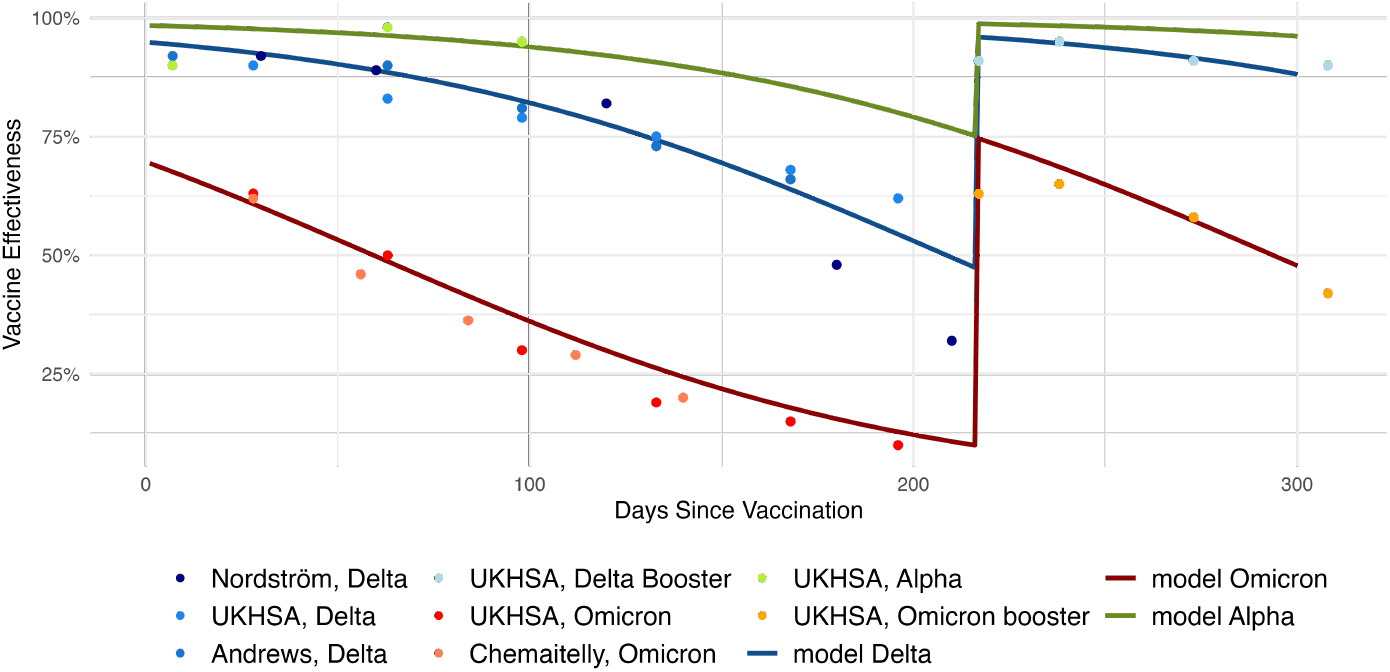
Calibration results. Dots were taken from the literature, lines are the fitted curves. On day 210 the agent receive a booster dose, which increases their level of protection.

### 4.3 Antibody Models for Epidemiological Predictive Modelling of COVID-19: A Literature Search

The previous section described our approach for modelling antibody levels in an epidemiological context. In this section, we present an overview of similar approaches that exist in the literature, as of July 2022. We compiled a list of all models that have been listed in one of the following resources: (a) the *Covid-19 Forecast Hub*^2^ ([Cra+21]), (b) the *European Covid-19 Forecast Hub*^3^ ([She+22a; She+22b]), and (c) the *European Covid-19 Scenario Hub*^4^ ([Eur22]). The final list contains 90 models. Additionally to the 86 models from the three resources, we also included four more models that we found through a PubMed literature search. The full list can be found in Appendix A. To get the relevant information for the individual models, we went to the respective websites, and analyzed connected publications and available source codes (e.g. from GitHub). We were in particular interested in models that either (1) related antibodies and protection against infection and integrated this into their model or (2) acknowledged and integrated into their model the waning of protection against infection (after vaccination and/or infection).

From this literature review, we conclude that, apart from Covasim ([Ker+21; Coh+21]), whose influence on our model has been discussed in Sec. 4.2.1 and which can be found as model #1 in Appendix <u>A),</u> none of the reviewed models explicitly integrate antibody levels as part of their infection sub-model. This is also due to the fact that many models focused on the prediction of hospitalization numbers and thus do not need to explicitly model individual antibody levels. However, some approaches seem to have integrated some kind of vaccination or antibody sub-model, but no detailed description is available. This includes (see table for details): IHME, UC3M-EpiGraph, ECDC-CM ONE, and SIMID-SCM.

## 5 Data and code availability

The data used in this study are all available from public resources that have been appropriately cited within the manuscript. Any additional information required to run the model is available from the lead contact upon request.

## Acknowledgements

The work on the paper was funded by the Ministry of research and education (BMBF) Germany (grants number 031L0300D, 031L0302A) and under Germany’s Excellence Strategy—MATH+ : The Berlin Mathematics Research Center (EXC-2046/1)—project no. 390685689 (subproject EF4-13). Additional resources have been provided by TU Berlin und Zuse Institute Berlin.

We thank Dr. Johannes Nießen, Prof. Dr. Dr. Alex Lechleuthner, Dr. Julia Hurraß from the Cologne Health Department for many insightful discussions.

## Author contribution

S.M., K.N., T.C. conceptualized the idea; S.M., S.P., J.R. curated the data; S.M., S.P., J.R. performed the experiments and S.M. did the formal analysis. S.P. did the literature review and S.M., S.P., J.R., K.N., T.C. wrote the original draft. All authors read, reviewed and approved the manuscript.

## Appendices

## A Appendix

### A.1 Literature Review

EMI: (YES, NO, N/A) Explicit Modelling of immunisation events (distinguish between infection, immunisation etc.) AND/OR of waning AND/OR of a relation between antibodies and protection against infection. If the model description solely mentions any of the criterion above, but provides no details, we still set EMI = N/A. If the same model was, under a slightly different name, part of multiple of our sources, we put down all names separated by a “/”.

**Table 2:**
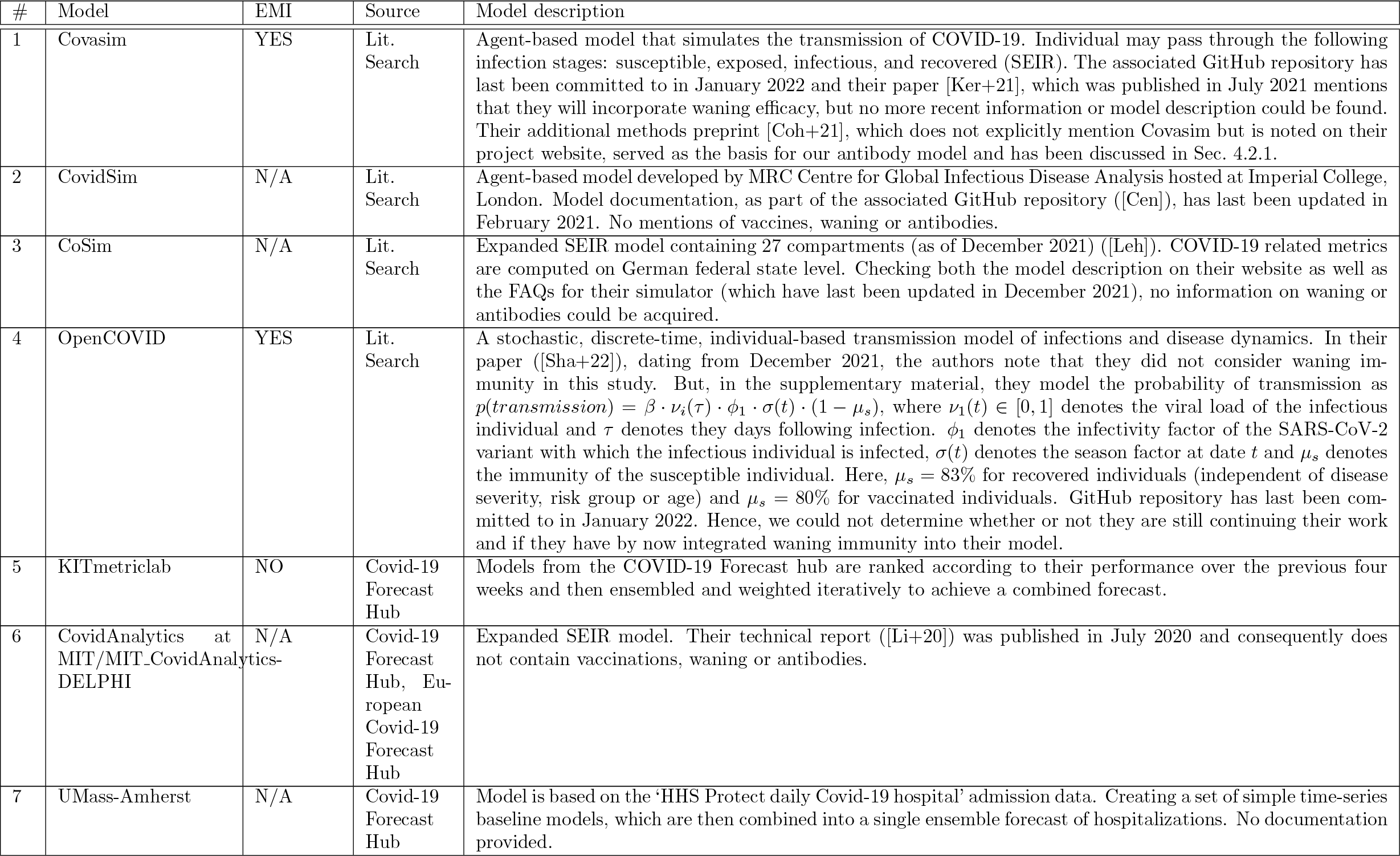
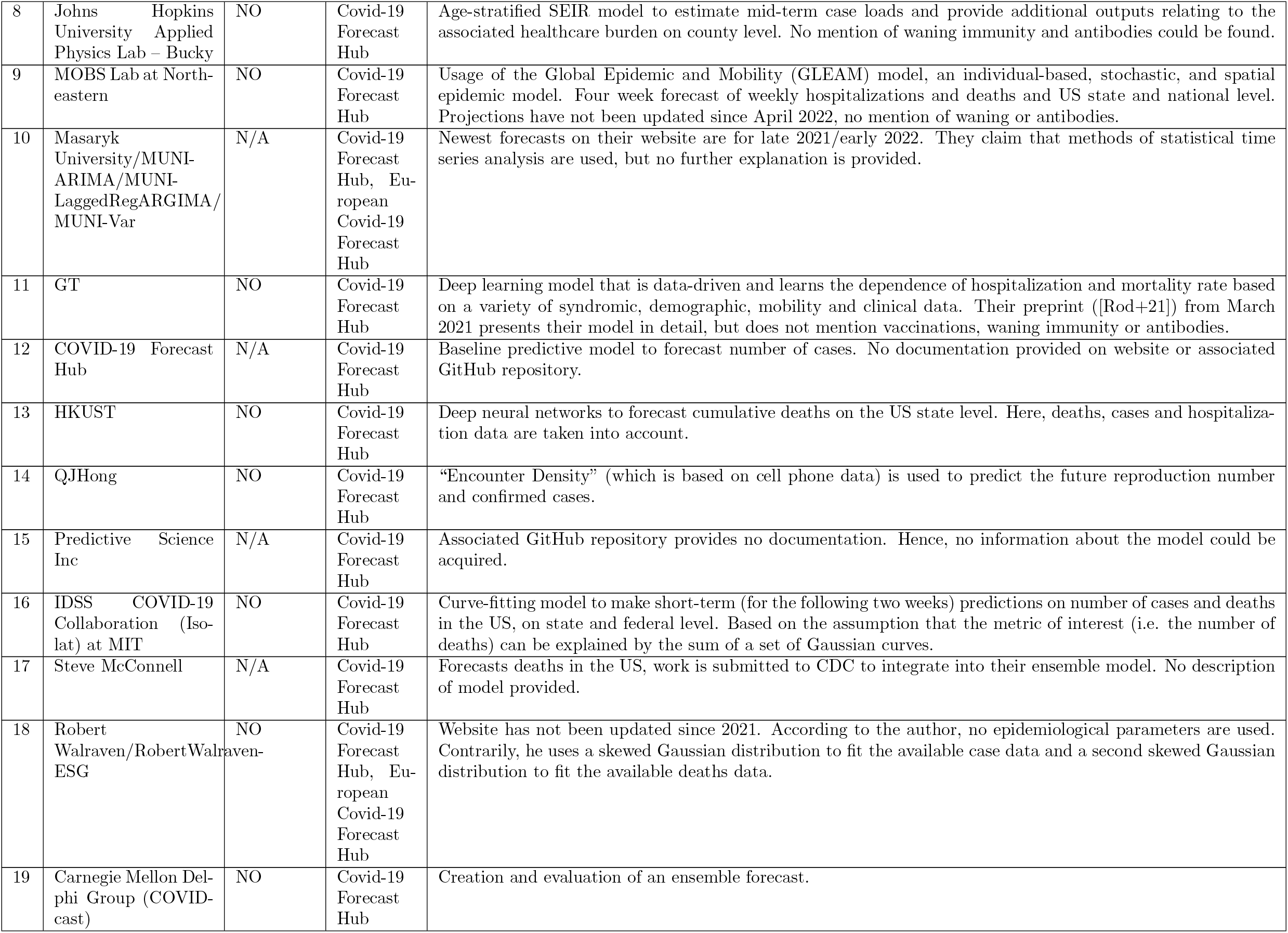
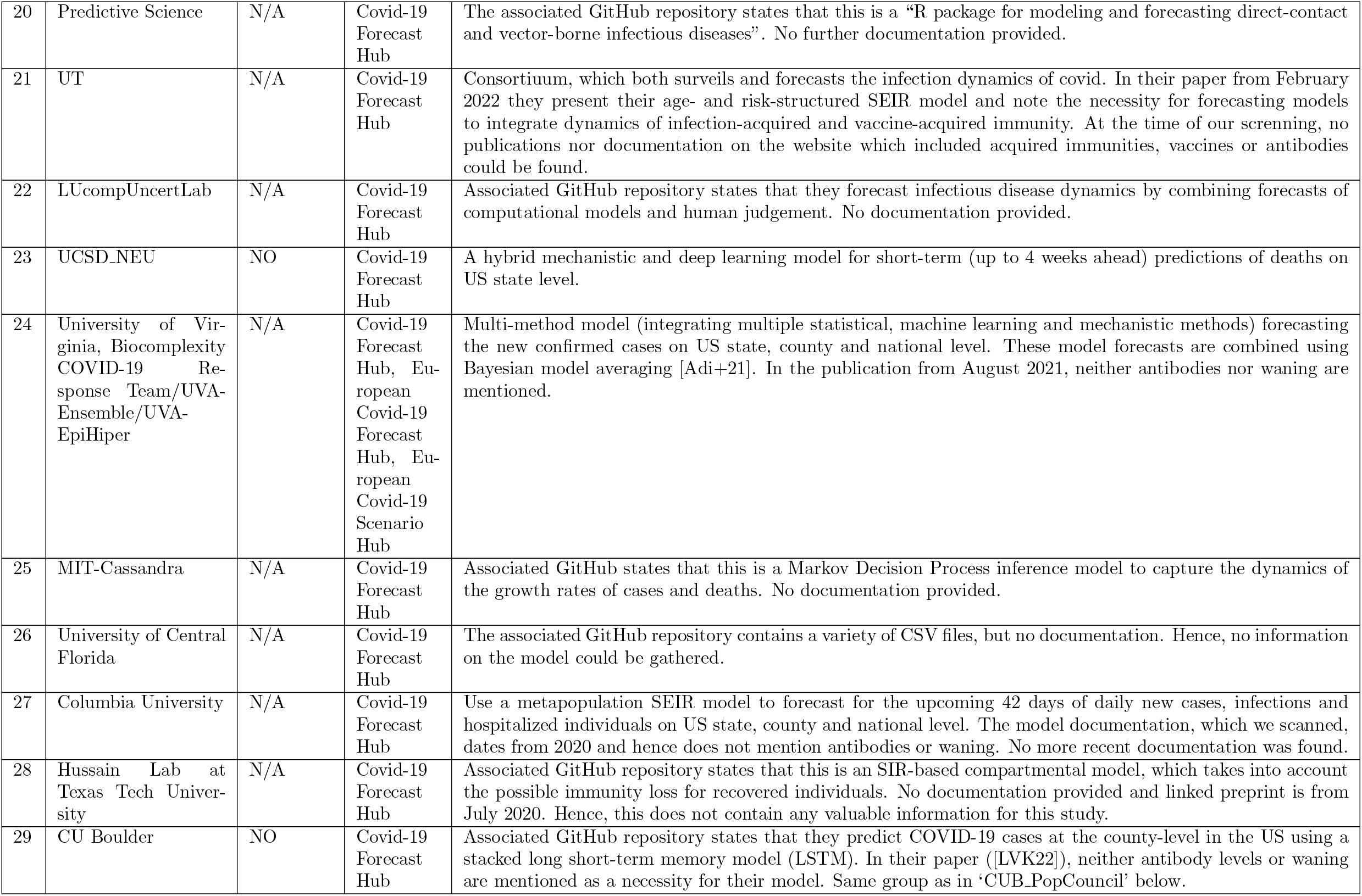
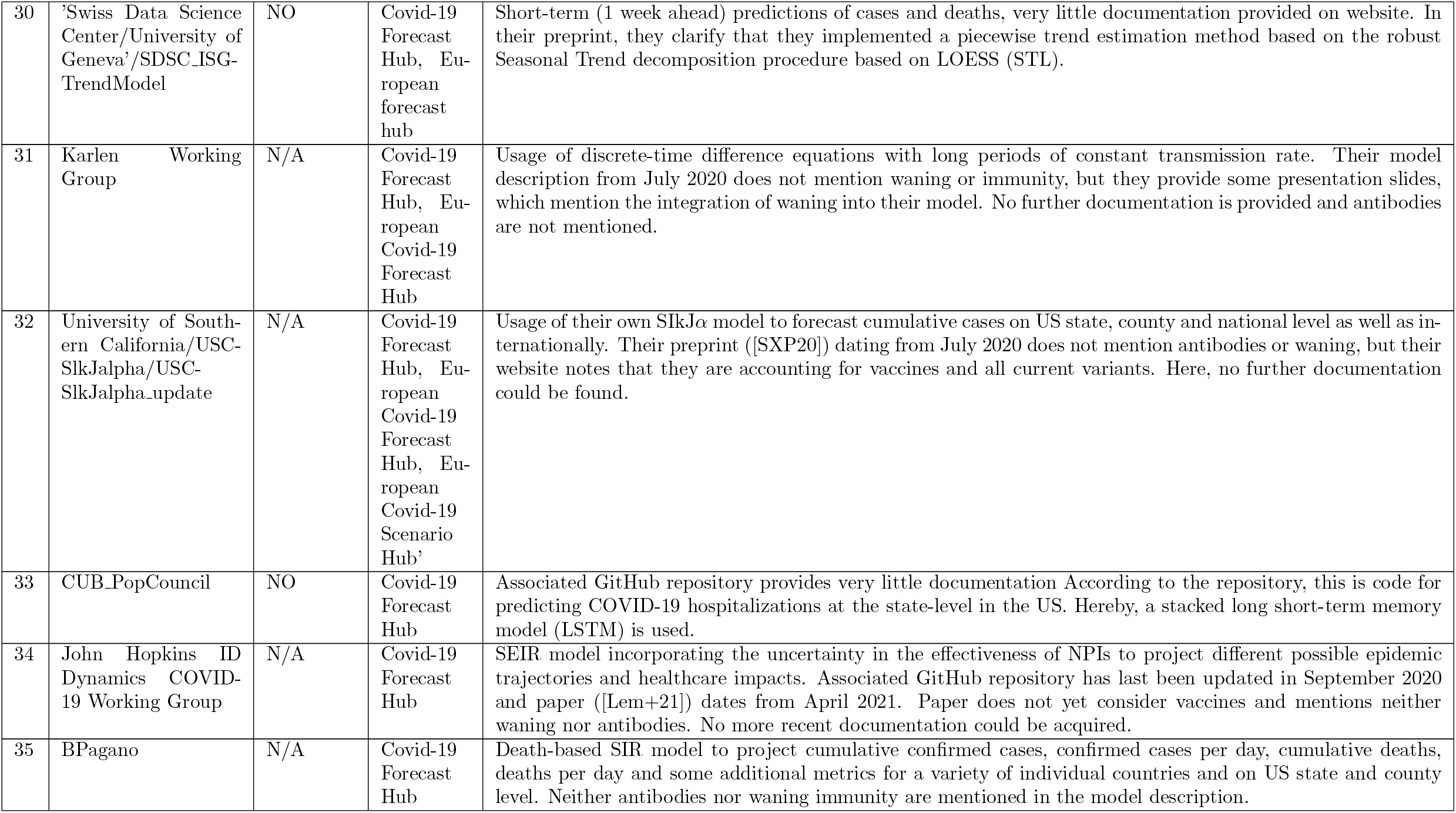
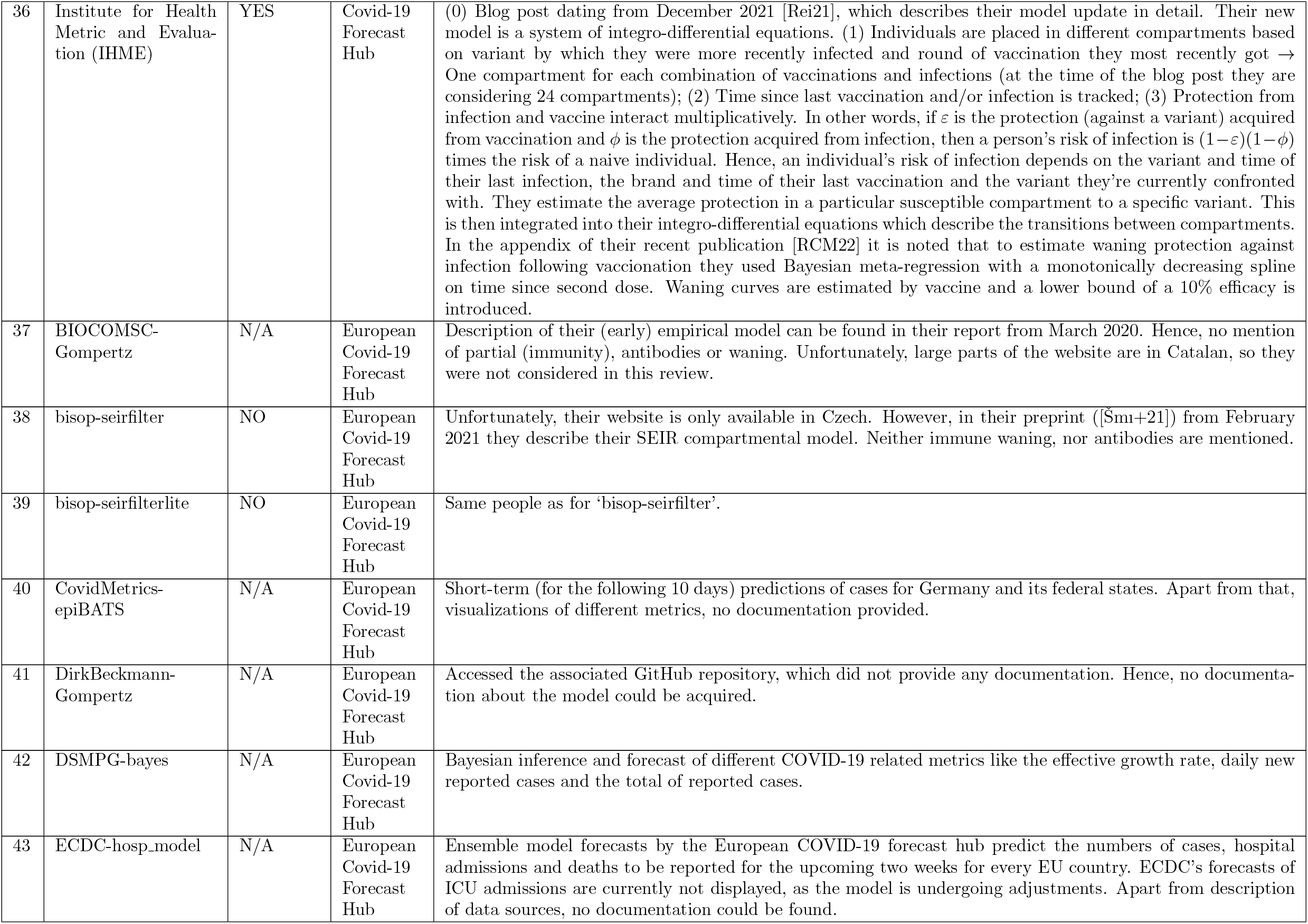
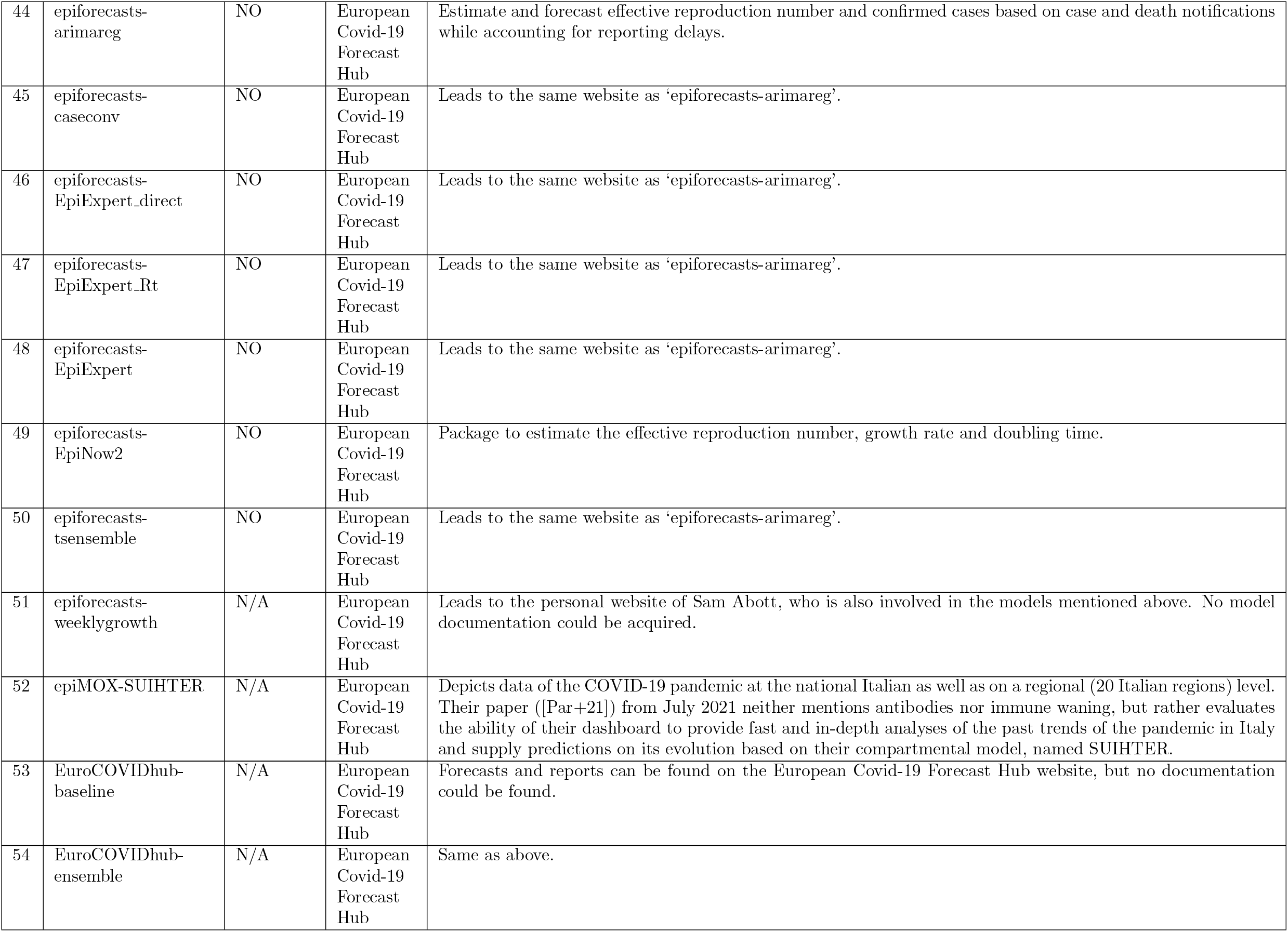
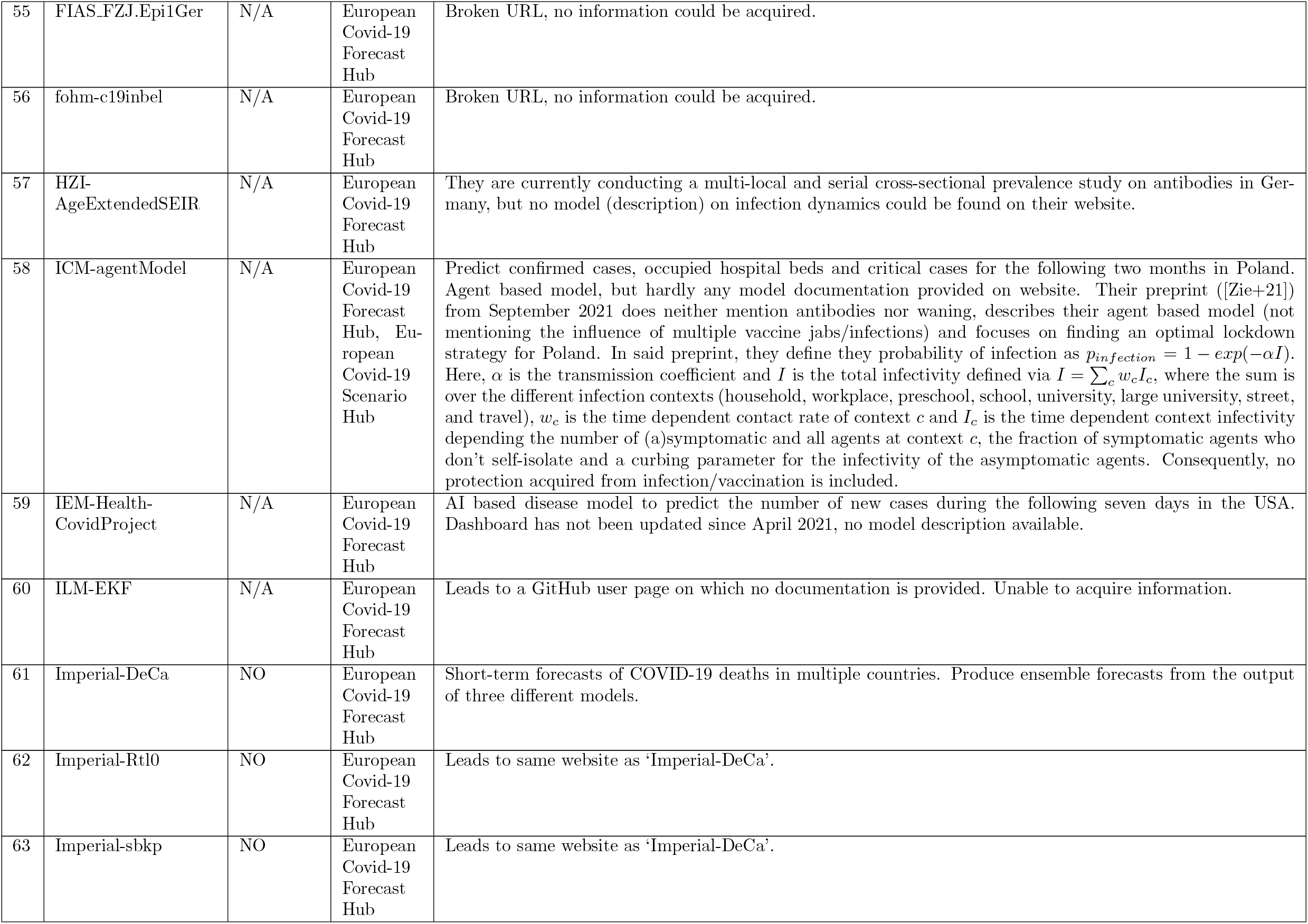
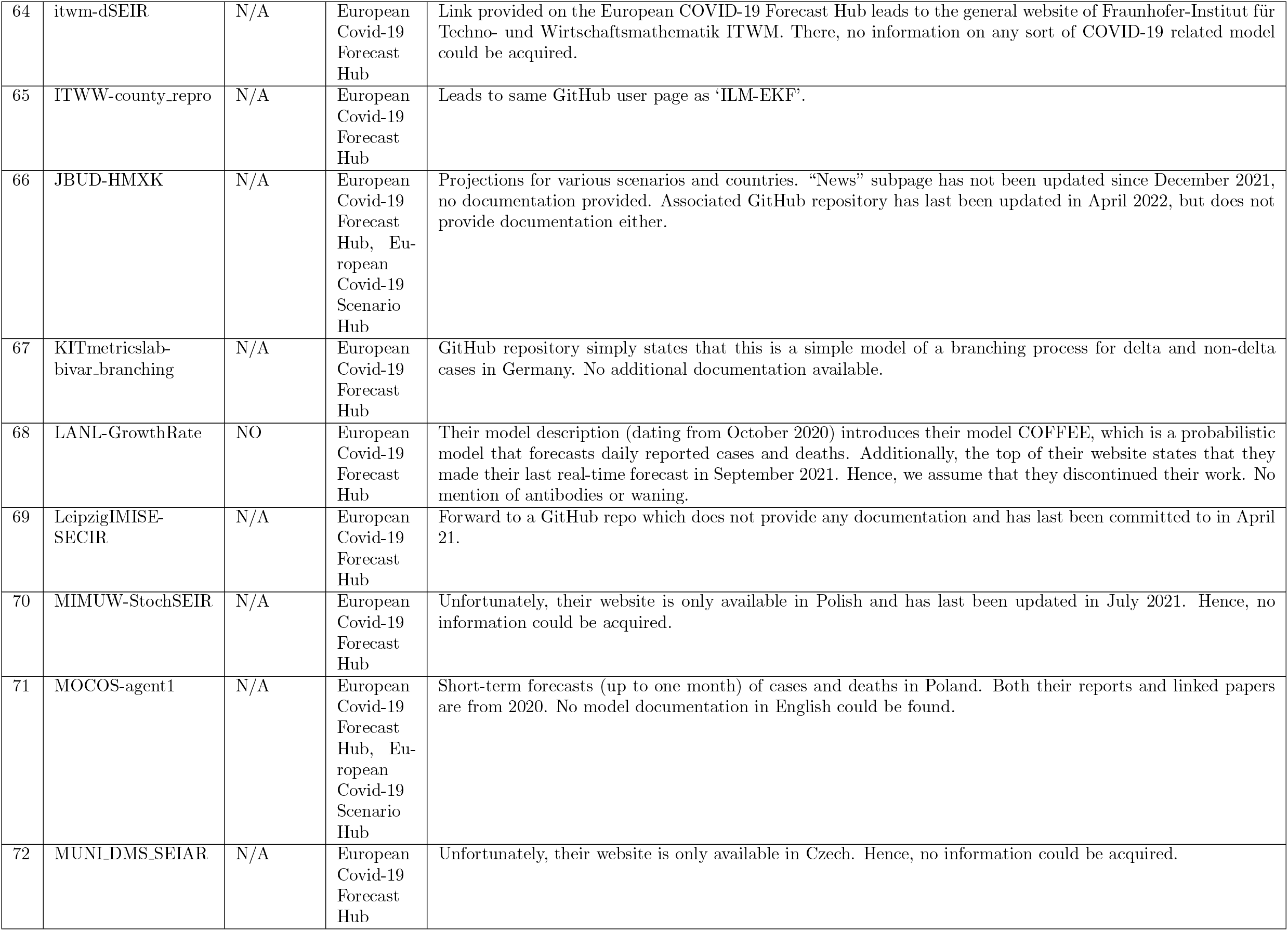
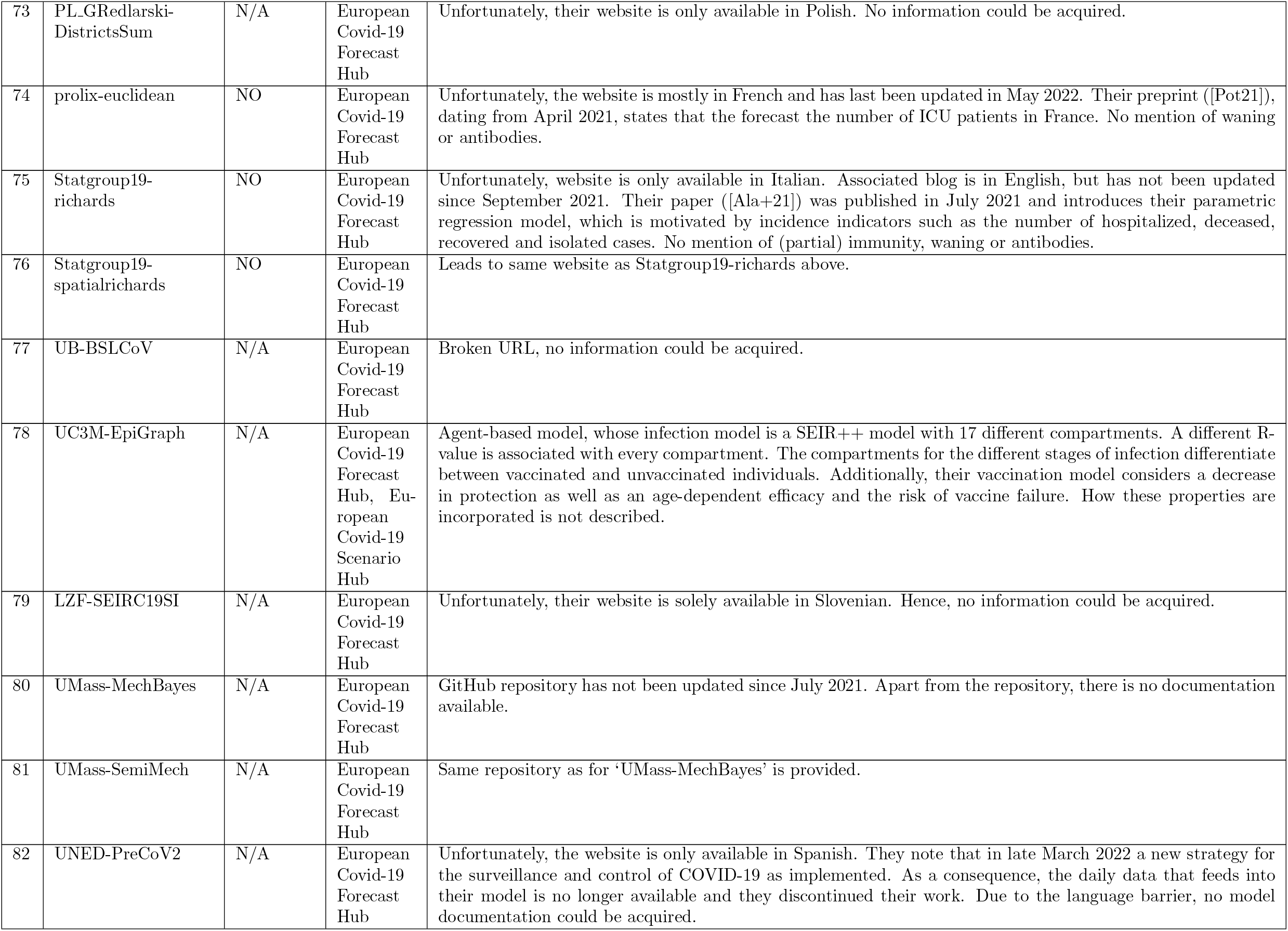
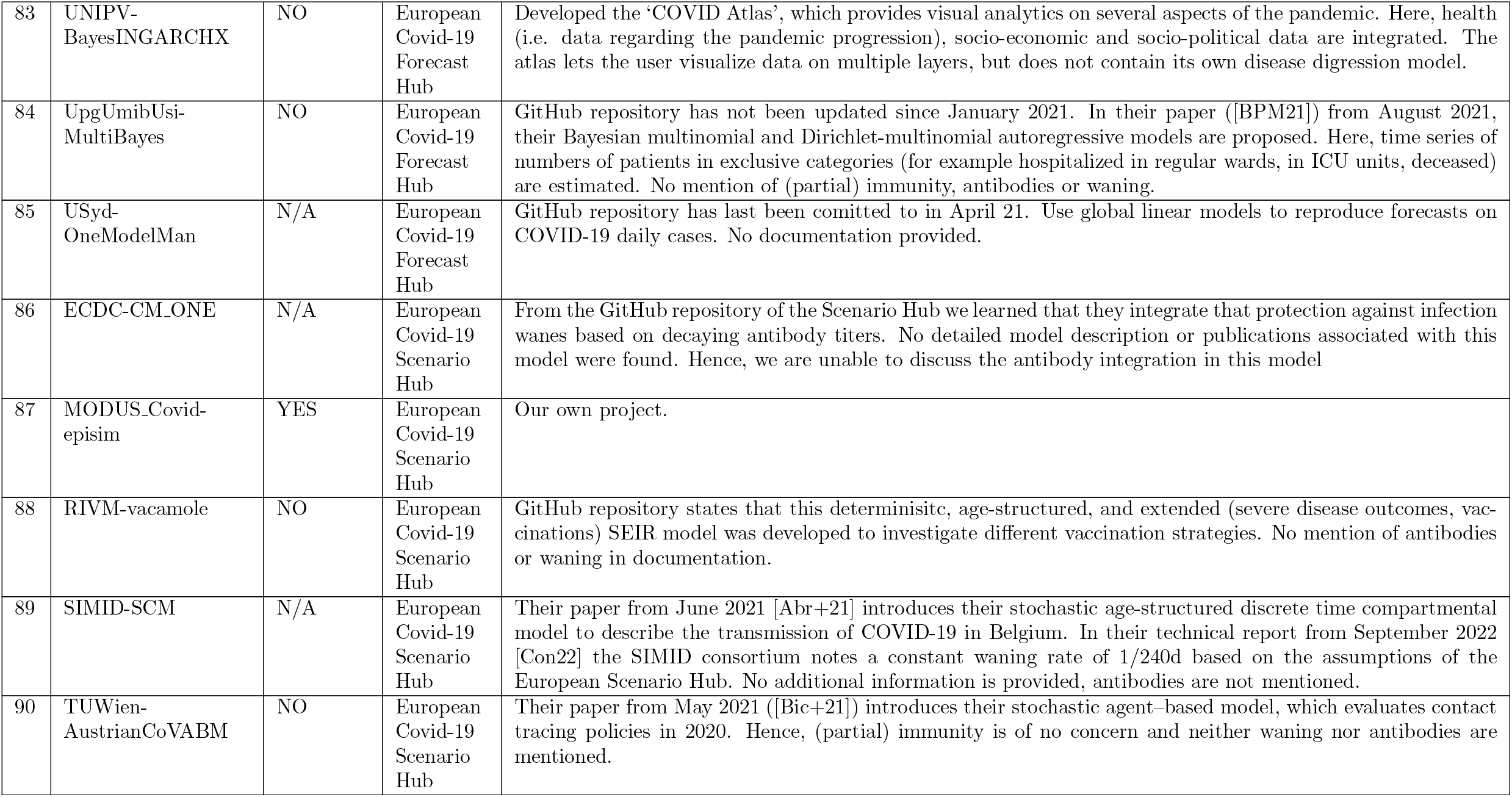
Models, which were discovered through (a) the Covid-19 Forecast Hub, (b) the European Covid-19 Forecast Hub, (c) the European Covid-19 Scenario Hub, or (d) a literature search. (a) and (b) were accessed on 07/21/22, while (c) was accessed on 08/31/22.

## Footnotes

1 A typical value for Θ in the model of [Mül+21] is of the order of 10^*−*5^. At the same time, without protection (e.g. masks) *d* = *τ/*(*rs · ae*), where *τ* is the time of exposure in seconds, *rs* is room size in *m*^2^, and *ae* is the air exchange rate per hour. Assume *rs* = 20*m*^2^ and *ae* = 0.5*/h*, typical values for a two-person office or a smallish living room, and *τ* = 1000*sec* of exposure time. These values result in Θ *· d* = 0.001.

2 From the community sub-section, as of 07/21/22.

3 From the community sub-section, as of 07/21/22.

4 From the models sub-section, as of 08/31/22.

